# TbTim20 facilitates protein import at a low membrane potential in trypanosomes lacking the mitochondrial genome

**DOI:** 10.1101/2025.04.25.650624

**Authors:** Corinne von Känel, Salome Aeschlimann, Michaela Husová, Silke Oeljeklaus, Philip Stettler, Achim Schnaufer, Bettina Warscheid, Alena Zíková, André Schneider

## Abstract

Protein import across the mitochondrial inner membrane typically depends on two protein translocases of the inner membrane (TIM) complexes and the membrane potential. The protozoan parasite *Trypanosoma brucei*, however, has a single, divergent TIM complex. Unlike other trypanosomal TIM subunits, TbTim20 is neither essential for normal growth of insect nor bloodstream forms of *T. brucei*, leaving its role uncertain. Specific mutations in the γ-subunit of the F_1_F_O_-ATPase, such as γL262P, permit bloodstream form trypanosomes to grow without mitochondrial DNA (kinetoplast or kDNA). Here we show that RNAi-mediated depletion of TbTim20 inhibits growth of this cell line, but only if it lacks the kDNA. Titration of mitochondrial uncouplers and direct membrane potential measurements reveal that TbTim20 becomes more critical as the membrane potential decreases across all tested cell lines. Proteomic analysis of the uninduced and induced γL262P TbTim20-RNAi cell line, which lacks kDNA and exhibits the lowest membrane potential, shows depletion of a subset of imported proteins. This subset includes ATPase subunits, suggesting a mechanism by which TbTim20-silenced cell lines become more sensitive to uncouplers. Thus, we propose that TbTim20 supports import of a subset of proteins whose import is hypersensitive to a low membrane potential.

## Introduction

Mitochondrial protein import has been studied since many decades and the machineries and mechanisms of the process are well understood (1, 2, 3). Historically, essentially all experimental studies of protein import have been restricted to yeast and mammals, which belong to the same eukaryotic supergroup of the Opisthokonts (4).

In the last decade, studies in the parasitic protozoan *Trypanosoma brucei*, a member of the Discoba group, revealed that its mitochondrial protein translocases are highly diverged when compared to Opisthokonts (5, 6, 7). The most extreme differences are found in the translocases of the inner membrane (TIM). Most eukaryotes have two TIM complexes, TIM23 and TIM22 (8, 9, 10, 11), that are specialized to import presequence-containing proteins and mitochondrial carrier proteins with multiple transmembrane domains, respectively. Trypanosomes, however, have a single bifunctional TIM complex that, with minor compositional variations, imports both types of substrates (12). Moreover, of all six integral membrane subunits of the trypanosomal TIM complex (12), the only one showing homology to any subunit of the TIM23 and TIM22 complexes of other eukaryotes is TbTim17 (13), an orthologue of Tim22 (14).

The energy requirements for mitochondrial protein import have mainly been studied in yeast and mammals (15). It has been shown that the inner membrane (IM) potential (ΔΨ) is required for electrophoretic translocation of the basic presequences across the TIM23 complex (16). The subsequent translocation of the mature part of import substrates depends on ATP and is mediated by mitochondrial Hsp70 (mHsp70), the core component of the presequence translocase-associated motor (PAM) (10, 17). Also in *T. brucei*, the ΔΨ is required to import presequence-containing substrates (18) and a modified PAM mediates ATP-dependent protein translocation (18, 19, 20, 21). Substrates of the yeast TIM22 complex do not require ATP but depend on the ΔΨ for both substrate docking and membrane insertion (22). In *T. brucei*, insertion of mitochondrial carrier proteins by the TIM complex also appears to depend on the ΔΨ, however, what precise role it plays is unknown (23).

Import substrates have a variable number of positive charges in their presequences and the mature parts of the proteins. Thus, conditions influencing the ΔΨ can affect protein import in a substrate-specific manner (24). While it is not fully understood how mitochondria guarantee import of proteins under both high and low ΔΨ conditions, this ability is likely important to maintain proper proteostasis.

The life cycle of *T. brucei* involves a tsetse fly vector and a mammalian host (25). The procyclic form (PCF) in the fly gut and the disease-causing long slender bloodstream form (BSF) have very different energy metabolisms (26, 27). Both life cycle stages can be grown in culture and therefore are excellent systems to study the effects this has on mitochondrial protein import.

PCFs produce most ATP by oxidative phosphorylation (OXPHOS) (28) (Fig. S1A), which requires proteins encoded on the mitochondrial genome, termed kinetoplast DNA (kDNA) in trypanosomes. BSFs produce most, if not all, ATP by glycolysis (29). Their mitochondria lack OXPHOS complexes, except for the F_o_F_1_-ATPase, which consumes ATP to generate the ΔΨ required for protein import and metabolite transport (30, 31) (Fig. S1B). Therefore, although BSFs are not capable of OXPHOS, they still depend on the kDNA, because subunit *a* of the F_o_-ATPase (32), ribosomal RNAs and two mitoribosomal proteins (33) are mitochondrially-encoded.

*T. b. equiperdum* and *T. b. evansi*, two subspecies of *T. brucei*, which lack most of or even all kDNA, cannot grow in the fly but are able to infect horses and a variety of mammals, respectively (34). Growth of the BSFs of these subspecies is possible due to specific mutations in their nucleus-encoded γ-subunit of the F_o_-ATPase, which compensates for the lack of kDNA-encoded subunit *a* (30, 35, 36, 37). The best evidence for this scenario is that a transgenic BSF *T. brucei* strain carrying a L262P mutation in one allele of the γ-subunit of the F_1_-ATPase, termed γL262P strain, grows as well as wild-type BSF even in the absence of kDNA, at least *in vitro* in rich medium (36) (Fig. S1C, S1D). In absence of kDNA and subunit *a*, the F_1_ and F_o_ parts of the ATPase are uncoupled (Fig. S1D). The γL262P mutation likely allows efficient ATP-hydrolysis by the uncoupled F_1_-part to enhance electrogenic exchange of cytosolic ATP^4−^ for an excess of mitochondrial ADP^3−^ by the ADP/ATP carrier, creating a ΔΨ of sufficient magnitude to allow cell growth (36). It is important to note that the concept of ΔΨ generation in the absence of the mitochondrial genome described above is not unique to trypanosomes, but was first discovered in petite mutants of yeast (38).

Here, we have analyzed the function of TbTim20, a subunit of the single trypanosomal TIM complex (39). We show that the protein is essential for normal growth of the γL262P strain lacking the kDNA, but not for kDNA-containing PCFs and BSFs when grown under standard conditions. Our results support a scenario that TbTim20 assists import of a subset of proteins, whose import is sensitive to a low ΔΨ.

## Results

### TbTim20 is an intermembrane space-localized TIM subunit

We have previously identified TbTim20 and TbTim15 as novel kinetoplastid-specific subunits of the trypanosomal TIM complex (39). Unlike the soluble intermembrane space (IMS)-localized TbTim15, RNAi-mediated depletion of TbTim20 did not affect growth of PCF cells, raising the question what its function might be.

Combining stable isotope labeling by amino acids in cell culture (SILAC) with co-immunoprecipitation (CoIP) and quantitative mass spectrometric analyses in PCF *T. brucei* using either TbTim20 or TbTim15 as baits (Fig. 1A, 1B) recovered TbTim20, TbTim15 and the membrane integral IM protein TimRhom I as the three most enriched proteins (including the baits) followed by essentially all other known TIM (12) and PAM subunits (Table S1, S2) (19). These results suggest a close association of TbTim20, TbTim15 and TimRhom I. Intriguingly, TbTim20 and TimRhom I are selectively associated with the active presequence but not the carrier translocase (12).

**Figure 1.**
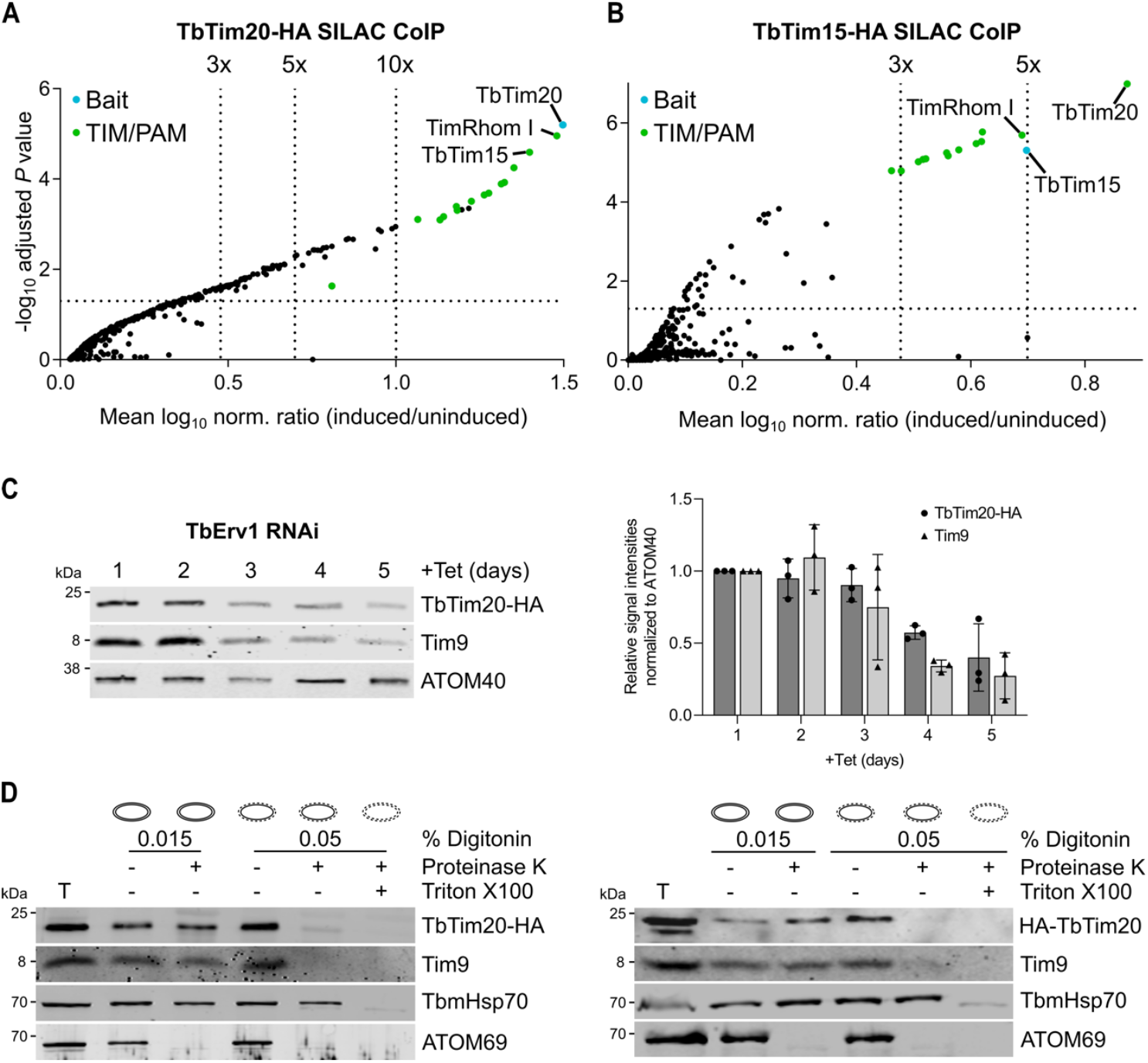
TbTim20 is an intermembrane space-localized TIM subunit. Visualization of proteins detected in SILAC-based quantitative mass spectrometry analysis of TbTim15-HA **(A)** and TbTim20-HA **(B)** CoIPs. Differentially labelled uninduced and induced cells were mixed and subjected to CoIP. The vertical dotted lines indicate the specified enrichment factors. The horizontal dotted lines indicate an adjusted *P* value of 0.05. The CoIP baits are highlighted in blue and TIM and PAM components in green. For numerical data see Table S1 and S2. **(C)** Left panel, immunoblot of steady-state levels of TbTim20-HA in TbErv1 RNAi background over five days of induction. Tim9 and ATOM40 were used as markers for mitochondrial intermembrane space and outer membrane proteins, respectively. Right panel, densitometric quantification of immunoblot signals of TbTim20-HA and Tim9 normalized to ATOM40 of triplicate experiments shown on left panel. Error bars indicate the standard deviation **(D)** Total cells (T) expressing either C- or N-terminally HA-tagged TbTim20 (TbTim20-HA, HA-TbTim20) were extracted with 0.015% digitonin to obtain mitochondria-enriched fractions. Treatment of these fractions with 0.05% digitonin allowed the generation of mitoplasts. Mitochondria-enriched fractions and mitoplasts were treated with 0.05 mg/ml proteinase K and/or 1% Triton X100 as indicated. The resulting immunoblots were probed with anti-tag antibodies. Tim9, TbmHsp70 and ATOM69 served as controls for mitochondrial intermembrane space, matrix, and outer membrane proteins, respectively.

Erv1 is a conserved protein that mediates import of IMS-localized substrates. Thus, the fact that TbTim20, similar to TbTim15 (39), was depleted upon TbErv1 RNAi indicates its localization in the IMS (40) (Fig. 1C). Protease protection assays using mitochondria and mitoplasts (i.e. isolated mitochondria with disrupted outer membrane) confirmed that both the N- and the C-terminus of TbTim20 are exposed to the IMS (Fig. 1D). Although TbTim20 contains no predicted transmembrane domains, it is tightly associated with the IM, as shown by its recovery in the pellet of an alkaline carbonate extraction (39). The structure of TbTim20 predicted by AlphaFold2 (41) consists of a bundle of α-helices (Fig. S2).

### TbTim20 is essential in kDNA-lacking ΨL262P cells

RNAi-mediated depletion of TbTim20 has no impact on growth of PCF (39), BSF wild-type cells (New York single marker, NYsm) (Fig. 2A), or the BSF kDNA-containing γL262P(+kDNA) strain (Fig. 2B, left panel). However, in the γL262P(-kDNA) TbTim20 RNAi cell line, where the kDNA was removed by ethidium bromide treatment, TbTim20 depletion led to a growth arrest (Fig. 2B, right panel). Note the γL262P mutation does not disrupt association of TbTim20 with the TIM complex (Fig. S3).

**Figure 2.**
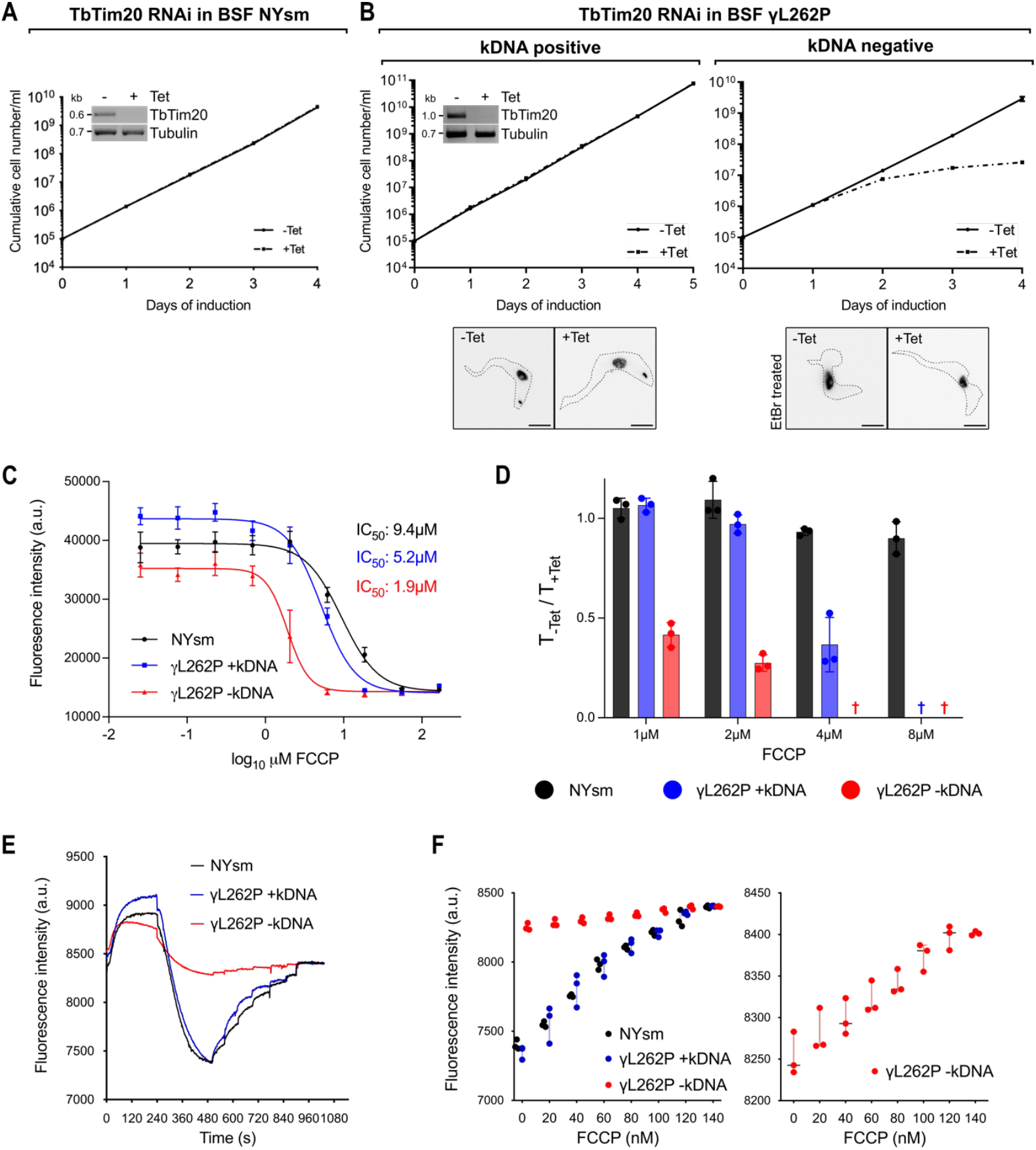
TbTim20 is essential for normal growth of BSFs with a decreased ΔΨ. **(A)** Growth curve of uninduced and induced (-/+Tet) TbTim20 RNAi BSF NYsm cells. Error bars are too small to be visible and correspond to the standard deviation (SD) (n=3). Inset shows the RT-PCR product of the TbTim20 mRNA in uninduced or two days induced cells. RT-PCR product of Tubulin mRNA serves as loading control. **(B)** Growth curves of uninduced and induced (-/+Tet) TbTim20-RNAi γL262P cell lines containing or lacking kDNA (+/-kDNA). SDs (n=3) are indicated. Inset on the left as in (A). The microscopy pictures of DAPI-stained cells in the bottom panels confirm the presence or absence of kDNA in the corresponding TbTim20-RNAi γL262P cell lines (scale bar: 5 μm). To remove the kDNA cells were grown in medium containing 10 nM ethidium bromide (EtBr) prior to tetracycline induction. **(C)** Alamar blue cell viability assays to determine the FCCP sensitivity of the indicated cell lines. Means and SD (n=3) are indicated. **(D**) The means and SDs of triplicate experiments showing the ratios between the generation times of uninduced and TbTim20-depleted normal BSF cells (NYsm), ΨL262P(+kDNA), and ΨL262P(-kDNA) cells in the presence of the indicated concentration of FCCP are depicted. Generation times were measured between two to three days after tetracycline-induced TbTim20 depletion. A cross indicates that only dead cells or cell debris were observed. Means and SD (n=3) are indicated. **(E, F)** ΔΨ measurement in the indicated permeabilized cell lines. A decrease in safranine O fluorescence is indicative of the formation of the ΔΨ. For each cell line a representative trace of three independent experiments is shown. Panels (E) and (F) depict dependence of fluorescence intensity versus time or FCCP concentration, respectively. The right panel in (F) shows fluorescence intensity of γL262P(-kDNA) cells with an expanded y-axis.

### kDNA-lacking γL262P cells are hypersensitive to a protonophore

In both wild-type BSFs (NYsm) and heterozygous γL262P(+/-kDNA) cells, ATP hydrolysis by the ATPase supports the generation of the ΔΨ (Fig. S1B, S1C, S1D). F_1_ F_o_-ATPase can directly generate the ΔΨ via ATP-hydrolysis-powered proton pumping (Fig. S1B, S1C). The kDNA encoded subunit *a* forms the proton-entry and -exit channel. Thus, in the absence of kDNA, proton translocation by the ATPase is no longer possible because the F_1_ and F_o_ parts of the ATPase are functionally uncoupled, leaving ΔΨ generation to the ADP/ATP carrier, presumably supported by the γL262P-containing F_1_-ATPase (Fig. S1D) (42). As mitochondrial protein import across the IM depends on the ΔΨ, the selective requirement for TbTim20 in γL262P(-kDNA) cells might be due to a lower ΔΨ in these cells when compared to γL262P(+kDNA) cells.

Sensitivity to the protonophore carbonylcyanide p-trifluoromethoxyphenylhydrazone (FCCP) can serve as a proxy for the magnitude of the *in vivo* ΔΨ. Thus, we determined the half maximal inhibitory concentration (IC_50_) of FCCP for all three cell populations using the Alamar blue viability assay. Fig. 2C shows that wild-type BSF (NYsm) cells were most resistant to FCCP, whereas γL262P(+kDNA) cells showed a medium and γL262P(-kDNA) cells the highest sensitivity to the protonophore. These results suggest that, as proposed above, the *in vivo* ΔΨ is higher in γL262P(+kDNA) than in γL262P(-kDNA) cells. Moreover, wild-type BSFs (NYsm) appear to have a higher *in vivo* ΔΨ than γL262P(+kDNA) cells. FCCP sensitivity was also influenced by TbTim20. The ratio between the generation times of the uninduced and induced TbTim20 RNAi cell lines at the indicated FCCP concentration declined for γL262P(+kDNA) and γL262P(-kDNA) cells, whereas wild-type BSF (NYsm) cells were only marginally affected (Fig. 2D).

### Loss of kDNA reduces ΔΨ of γL262P cells

FCCP sensitivity of cells serves as a proxy for the ΔΨ, but it does not directly measure it. To assess to which extent mitochondria of the three cell lines can develop a ΔΨ, we used the fluorescent dye safranine O as a quantitative indicator of the ΔΨ. Addition of 1 mM ATP resulted in a strong and comparable decrease in safranine O fluorescence in permeabilized wild-type BSFs (NYsm) and γL262P(+kDNA) cells, indicative of the formation of the ΔΨ (Fig. 2E, black and blue curves). However, the ATP-induced decrease in fluorescence observed in γL262P(-kDNA) cells was greatly diminished under the same conditions (Fig. 2E, red curve). The subsequent stepwise addition of FCCP (20 nM each) resulted in a corresponding stepwise depolarization of the ΔΨ which was complete at 140 nM FCCP (Fig. 2F). Whereas mitochondria of wild-type BSFs (NYsm) and γL262P(+kDNA) cells exhibited a comparable ΔΨ, the magnitude of the ΔΨ produced by γL262P(-kDNA) cells was much reduced.

The results for the γL262P(+ or -kDNA) cells are in agreement with their observed *in vivo* FCCP sensitivity (Fig. 2C). However, wild-type BSFs (NYsm) and γL262P(+kDNA) have different *in vivo* FCCP sensitivities even though the ΔΨ they can produce in permeabilized cells under standard conditions is very similar (Fig. 2E). A possible explanation for this could be that the intracellular ATP levels are different in wild-type BSFs (NYsm) and γL262P(+kDNA) cells, suggesting that *in vivo* wild-type BSFs (NYsm) cells can generate a higher ΔΨ.

### TbTim20-depletion changes the mitochondrial proteome in kDNA-lacking γL262P cells

Fig. 3 shows proteomic comparisons between crude mitochondrial fractions of uninduced and induced γL262P(-kDNA) TbTim20 RNAi cells. This allows the global identification of import substrates that require TbTim20 for efficient import. As expected, TbTim20 was efficiently depleted (4.6-fold). A total of 1729 proteins were quantified, 363 of them were reported to be part of the BSF mitochondrial proteome based on a previous study (43) (Table S3). Of the remaining 1366 proteins five were predicted to contain a mitochondrial presequence resulting in a total number of 368 detected putative BSF mitochondrial proteins. Whereas of the 1729 quantified proteins 13% were depleted more than 1.5-fold, this fraction rose to 42% when only putative BSF mitochondrial proteins were considered, indicating that in the absence of kDNA a subset of mitochondrial proteins requires TbTim20 for efficient import. 94 of the 368 putative BSF mitochondrial proteins were predicted by TargetP (44) to contain N-terminal presequences, and 75% of these were depleted more than 1.5-fold (Fig. 3A, Table S3). These results suggest that the function of TbTim20 is linked to the presequence pathway. Intriguingly, nine subunits of the F_1_F_o_-ATPase and the ADP/ATP carrier were depleted more than 1.5-fold (Fig. 3A), potentially explaining why TbTim20-silenced cells have a lower ΔΨ (Fig. 2D).

**Figure 3.**
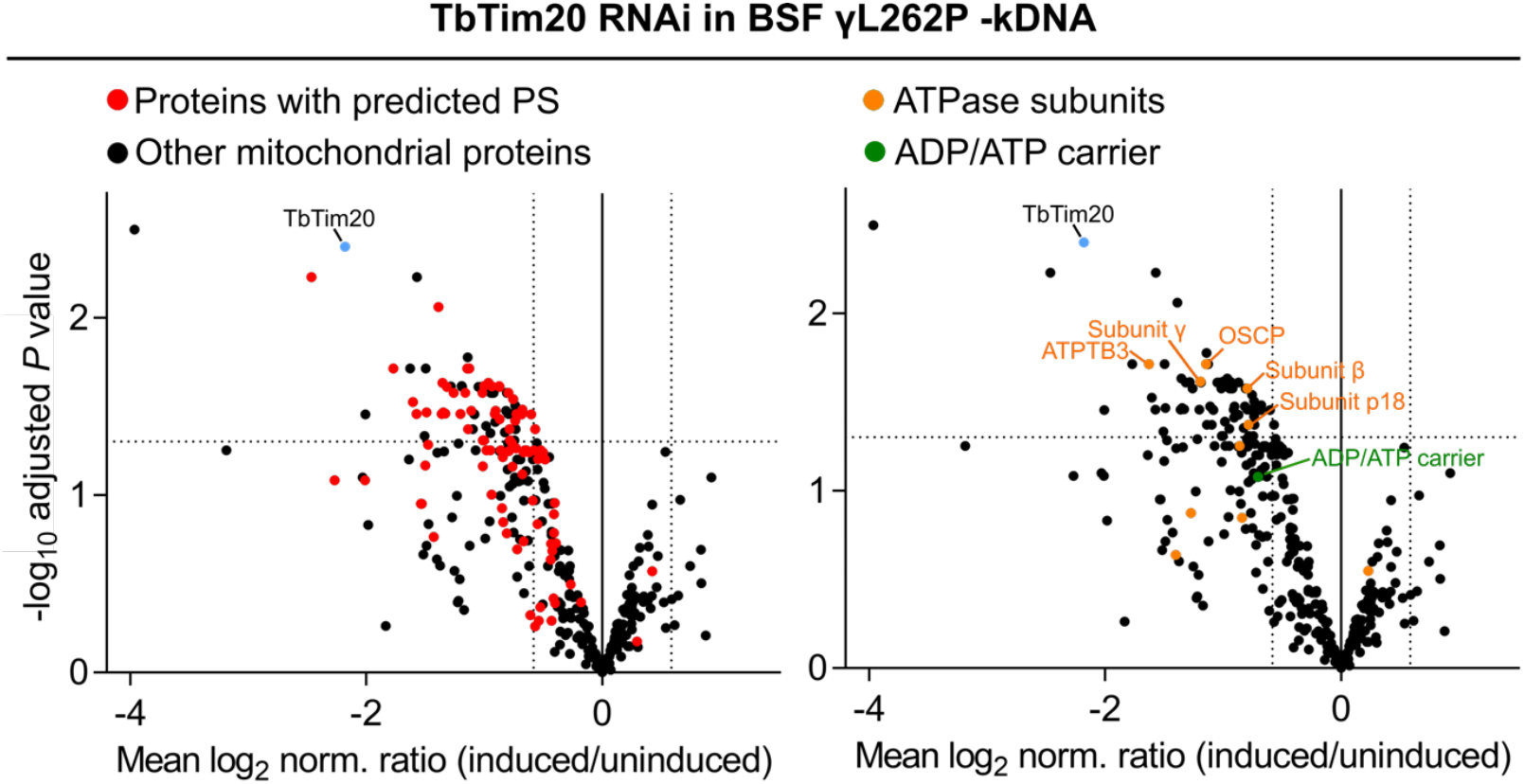
TbTim20-depletion changes the mitochondrial proteome in kDNA-lacking γL262P cells. Mitochondria-enriched fractions of uninduced and two days induced TbTim20 RNAi γL262P(-kDNA) were analysed by quantitative proteomics. To ablate the kDNA, cells were grown in medium containing 10 nM ethidium bromide starting three days before RNAi induction. Only putative BSF mitochondrial proteins are depicted. The mean log_2_ of normalized ratios (induced/uninduced) was plotted against the corresponding negative log_10_ of the adjusted *P* value (limma test). TbTim20 is highlighted in blue. Left: proteins with a predicted mitochondrial presequence are highlighted in red. Right: subunits of the ATPase and the ADP/ATP carrier are highlighted in orange and green, respectively. The horizontal dotted lines in both the volcano plots mark an adjusted *P* value of 0.05. The vertical dotted lines in both volcano plots indicate a +/-1.5-fold-change in protein abundance. For numerical data see Table S3.

However, no specific characteristics were identified that explain why at a low ΔΨ the depleted proteins might require TbTim20 for import. Neither the number of positive charges in the first 20 amino acids nor a higher isoelectric point (pI) correlated with the decrease of mitochondrial protein levels of the predicted presequence-containing proteins following TbTim20 depletion (Fig. S4).

### TbTim20 facilitates growth of PCFs in the presence of a protonophore

Under standard culture conditions RNAi-mediated knockdown of Tim20 resulted in only a marginal effect on growth of PCF trypanosomes after a prolonged cultivation. However, when a sublethal concentration (2.5 μM) of the uncoupler carbonyl cyanide m-chlorophenylhydrazon (CCCP) was added to the culture RNAi knockdown of Tim20 led to a strong growth retardation. Addition of CCCP decreases the ΔΨ mimicking the situation in BSF γL262P(-kDNA) cells.

## Discussion

Here, we show that TbTim20, a subunit of the single trypanosomal TIM complex, is required for efficient mitochondrial import of a subset of proteins in BSF *T. brucei* cells that lack kDNA (γL262P(-kDNA) cells) and thus is essential for normal growth of these cells. However, in PCF, wildtype BSF (NYsm), and γL262P(+kDNA) cells grown under standard conditions, which all have a higher mitochondrial ΔΨ than γL262P(-kDNA) cells, depletion of TbTim20 did not affect growth (Fig 2).

These results suggest that TbTim20 is required for efficient translocation of a subpopulation of proteins across the IM when the ΔΨ is low. This is reminiscent to the non-essential integral IM protein Pam17 of yeast. Pam17 supports import of the mature part of ΔΨ-sensitive matrix proteins after IM translocation of their presequences and before ATP-driven import of their mature parts (24).

TbTim20 and its close interaction partner TimRhom I (Fig. 1A) are associated with the active presequence but not with active carrier translocase (12, 39). Like yeast Pam17, RNAi-mediated depletion of TbTim20 causes a preferential depletion of presequence pathway substrates (Fig. 3). However, TbTim20 is an IMS-localized protein (Fig. 1C, 1D) and thus must act at a different step than the matrix-exposed Pam17 (45). As in the case of Pam17, which features of the precursor proteins make their import TbTim20-dependent at a low ΔΨ are unknown (Fig. S4). Thus, even though the TIM complexes of *T. brucei* and yeast are very different, both contain subunits that appear to support import of substrates that are hypersensitive to a low ΔΨ, albeit by different mechanisms.

TbTim20-depleted cells have an increased sensitivity to FCCP, suggesting a further decrease of the ΔΨ in all tested cell lines (Fig 1D). This is likely a secondary effect of the reduced import of nine subunits of F_1_F_o_-ATPase and/or possibly the ADP/ATP carrier, two of which have an experimentally determined presequence (46). The alternative explanation that TbTim20 is directly involved in maintaining the ΔΨ, seems unlikely. It would be difficult to understand how TbTim20, a subunit of the TIM complex localized in the IMS, could directly affect the ΔΨ.

Our study also indicates that *in vivo* the mitochondrion in wild-type BSF cells (NYsm) has a higher ΔΨ than the one in heterozygous γL262P(+kDNA) cells, even though when tested in permeabilized cells under high external ATP concentrations their ΔΨ is very similar (Fig. 2C, 2E). This suggests that the L262P mutation in the γ-subunit of the ATPase, even when the wild-type version is still expressed, is sufficient to cause a reduction of the *in vivo* ΔΨ possibly because it lowers cellular ATP levels when uncoupled.

If TbTim20 is not essential for normal growth of PCF and wild-type BSF cells, why has its gene been maintained during evolution? It is plausible to assume that the *in vivo* ΔΨ can fluctuate depending on nutrient availability and environmental changes. Thus, we propose that TbTim20 allowed the maintenance of efficient protein import during such periods in both wildtype PCF and in BSF cells. The fact that TbTim20 depletion results in a growth retardation in PCF cells when grown in the presence of sublethal concentration of CCCP is in line witch such a role in metabolic adaptation (Fig. 4B).

**Figure 4.**
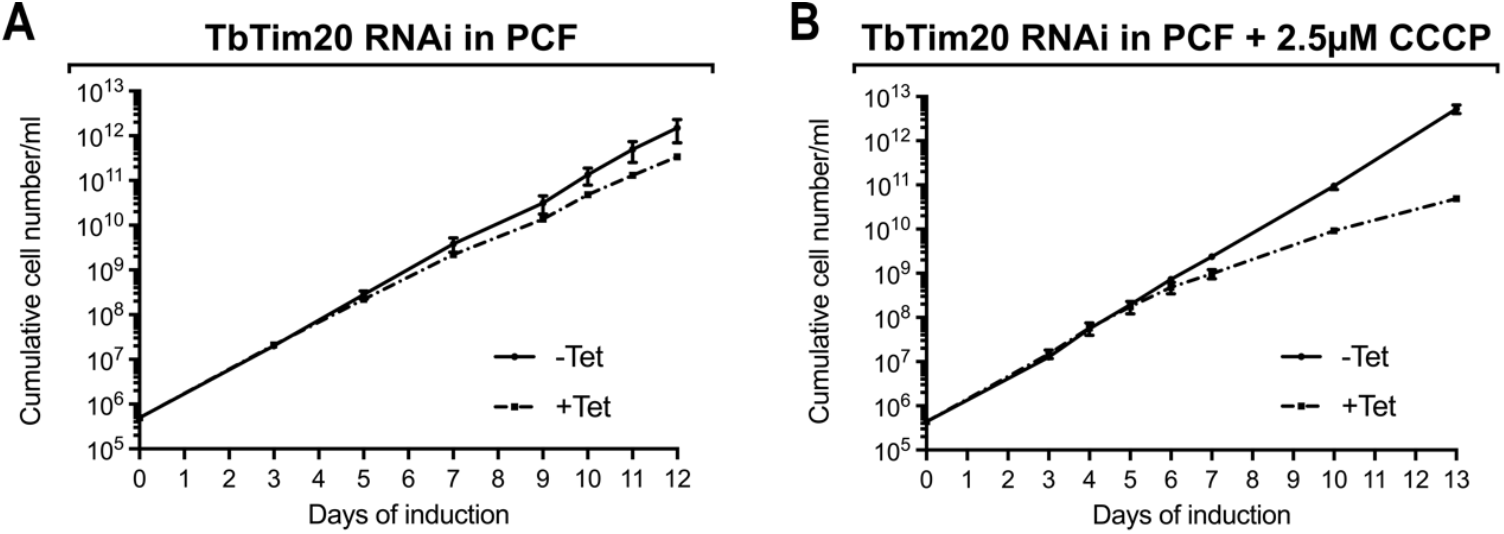
TbTim20 facilitates growth of PCF *T. brucei* in the presence of a protonophore. **(A)** Growth of uninduced and induced (-Tet/+Tet) PCF TbTim20 RNAi cells in standard medium. **(B)** As in (A) but growth was monitored in the presence of 2.5 μM of CCCP in standard medium. Error bars correspond to standard deviation (n=2).

## Material and methods

### Transgenic cell lines

Transgenic *T. brucei* cell lines are based on the PCF strain 29-13 (47) or the BSF wild-type strain (NYsm) (47) and γL262P (36). PCF cells were grown in SDM-79 or in SDM-80 (48) at 27°C, BSF parasites were grown in HMI9 (49) at 37°C, both supplemented with 10% (v/v) foetal calf serum (FCS). To induce loss of kDNA γL262P cells were cultured in presence of 10 nM ethidium bromide (EtBr) starting three days before the start of the respective experiment.

Plasmids for ectopic expression of C-terminal triple HA-tagged TbTim15 (Tb927.2.4445) and TbTim20 (Tb927.11.1620) have been described before (39). To generate a plasmid for ectopic expression of N-terminal triple HA-tagged TbTim20 the complete ORF was amplified by PCR and cloned into a modified pLew100 vector (47), which encodes a puromycin resistance gene and a N-terminal triple HA-tag (50).

The PCF TbTim20 RNAi cell line and the plasmids used to generate the TbTim20 BSF NYsm, γL262P RNAi cell line have been described before (39). Also the TbErv1 RNAi cell line has been described previously (40).

### Antibodies

Commercially available antibodies were: mouse anti-HA (Sigma-Aldrich, dilution immunoblot (IB): 1:5’000). Antibodies previously produced in our laboratory are: polyclonal rabbit anti-ATOM40 (51) (dilution IB: 1:10’000), polyclonal rabbit anti-ATOM69 (51) (dilution WB: 1:50), polyclonal rabbit anti-Tim9 (12) (dilution IB: 1:40), polyclonal rat anti-TbTim17 (12) (dilution IB: 1:300), polyclonal rabbit anti-TbPam27 (39) (dilution IB: 1:200) and polyclonal rabbit anti-TbTim15 (39) (dilution IB: 1:250-500). Monoclonal anti-TbmHsp70 (52) (dilution IB: 1:1’000) has described before. Secondary antibodies used were: goat anti-mouse IRDye 680LT conjugated (LI-COR Biosciences, dilution IB: 1:20’000), goat anti-rabbit IRDye 800CW conjugated (LI-COR Biosciences, dilution IB 1:20’000) and goat ani-rat IRDye 680LT conjugated (LI-COR biosciences, dilution IB 1:10’000).

### Digitonin extractions and protease protection assays

Generation of mitochondria-enriched digitonin pellets has been described in detail before (53). To generate mitoplasts, mitochondria-enriched digitonin pellets were incubated in a buffer containing 15 mM Tris base, 12 mM KH_2_PO_4_, 15 mM MgSO_4_, 0.5 M sorbitol pH 7.2 and 0.05% digitonin for 5 min at 4°C followed by centrifugation (6’700 g, 5 min, 4°C) yielding a pellet consisting of mitoplasts. Mitochondria-enriched pellets and mitoplasts were then resuspended in a buffer containing 20 mM Tris base, 15 mM KH_2_PO_4_, 20 mM MgSO_4_ and 0.6 M sorbitol pH 7.2 which was supplemented with 0.05 mg/ml proteinase K and/or 1% Triton X100, respectively.

The resulting samples were analyzed by SDS-PAGE and immunoblotting.

### Co-immunoprecipitation (CoIP)

The CoIP experiment shown in Fig. S3 was performed as follows. A mitochondria-enriched pellet from 1×10^8^ γL262P(+kDNA) cells expressing TbTim20-HA was generated by digitonin extraction as described before (53). This crude mitochondrial fraction was solubilized in 20 mM Tris-HCl (pH 7.4), 0.1 mM EDTA, 100 mM NaCl, 10% glycerol, 1× Protease Inhibitor mix (Roche, EDTA-free), and 1% (w/v) digitonin) for 15 min at 4°C. After centrifugation (20’000 g, 15 min, 4°C) the lysate was transferred to 50 μl HA bead slurry (anti-HA affinity matrix, Roche), which had been equilibrated in wash buffer containing 20 mM Tris-HCl (pH 7.4), 0.1 mM EDTA, 100 mM NaCl, 10% glycerol and 0.2% (w/v) digitonin. After incubation in an end-over-end shaker for at least one hour at 4°C, the supernatant containing the unbound proteins was removed. After washing the bead slurry three times with wash buffer, the bound proteins were eluted by boiling the resin for 5 min in 60 mM Tris-HCl (pH 6.8) containing 2% SDS. 5% of both the input and the unbound proteins, and 100% of the IP sample were analysed by SDS-PAGE and immunoblotting.

### Fluorescence microscopy

γL262P(+/-kDNA) cells were fixed with 4% paraformaldehyde in PBS, postfixed in cold methanol, and mounted using VectaShield containing 4′,6-diamidino-2-phenylindole (DAPI) (Vector Laboratories). Images were acquired by a DMI6000B microscope and a DFC360 FX monochrome camera (both Leica Microsystems).

### RNA extraction and northern blotting

Acid guanidinium thiocyanate-phenol-chloroform extraction to isolate total RNA from uninduced and induced (two days) RNAi cells was done as described before (54). To determine RNAi efficiency, the resulting RNA was used for RT-PCR or northern blotting as described previously (53).

### SILAC CoIP experiments

PCF cells inducibly expressing TbTim15-HA or TbTim20-HA were washed in PBS and resuspended in SDM-80 (55) containing 5.55 mM glucose, 10% dialyzed FCS (BioConcept, Switzerland) and either light (^12^C_6_/^14^N_χ_) or heavy (^13^C_6_/^15^N_χ_) isotopes of arginine (1.1 mM) and lysine (0.4 mM) (Eurisotope). To ensure complete labelling of all proteins with heavy amino acids, the cells were grown in SILAC medium for six to ten doubling times.

Digitonin-extracted mitochondria-enriched pellets of 4 x 10^8^ uninduced and one day induced cells were mixed and subjected to CoIP as described above. SILAC experiments were executed in three (TbTim20-HA) or four (TbTim15-HA) biological replicates including a label-switch and analyzed by liquid chromatography-mass spectrometry (LC-MS).

### Proteomic analysis of γL262P(-kDNA) TbTim20 RNAi cells

**γ**L262P TbTim20 RNAi cells were pretreated with 10 nM ethidium bromide (EtBr) starting three days before RNAi induction to induce loss of the kDNA. Digitonin-extracted, mitochondria-enriched pellets were generated from uninduced and two days induced BSF **γ**L262P(-kDNA) TbTim20-RNAi cells and analysed by quantitative LC-MS.

### Quantitative LC-MS and data analysis

Samples from SILAC CoIP and TbTim20 RNAi experiments were processed for LC-MS analysis as described before (40, 56) and measured using an Orbitrap Elite (TbTim20 CoIPs) or a Q Exactive Plus (TbTim15 CoIPs, TbTim20 RNAi experiments) mass spectrometer coupled to an UltiMate 3000 RSLCnano HPLC system (Thermo Fisher Scientific, Germany). Proteins were identified using MaxQuant/Andromeda (versions 1.6.3.4, 2.0.2.0 and 2.5.1.0 for TbTim20 CoIPs, TbTim15 CoIPs, or TbTim20 RNAi experiments, respectively) (57, 58) and a fasta file containing the protein sequences for *T. brucei* TREU927 downloaded from the TriTrypDB (https://tritrypdb.org; versions 8.1 and 55). MaxQuant was operated using default settings with the exception that only one unique peptide was required for protein identification.

For the analysis of SILAC data from CoIP experiments, Lys8/Arg10 were set as heavy labels and the option “requantify” was enabled. Quantification was based on ≥ one SILAC peptide pair.

Proteins significantly enriched in TbTim15 or TbTim20 complexes were identified using the rank sum method (56, 59, 60). The rank sum (i.e., the arithmetic mean of the ranks of a protein in all replicates based on normalized abundance ratios) was converted into FDR-controlled q-values. Effects of TbTim20 depletion on the mitochondrial proteome of BSF γL262P(-kDNA) cells were analyzed by label-free quantification using MaxQuant LFQ intensities. Only proteins present in at least 3/4 replicates of control samples (-Tet) were considered for further analysis. For each individual experiment, the median of LFQ intensities was shifted to the median of all medians. Next, LFQ intensities were transformed by variance stabilizing data normalization (vsn) (61).

Missing intensities were imputed by (i) the median value of the existing three replicates for control samples (-Tet) and (ii) minimum value imputation applying a downshift of 1.8 standard deviations and a width of 0.3 for tetracycline-treated samples (+Tet). Protein abundance ratios (+/-Tet) were calculated and q-values determined following the “linear models for microarray data” (limma) approach (62, 63) using the Benjamini-Hochberg method (64) for p-value correction.

Information about proteins identified and quantified in LC-MS experiments are available in Tables S1 - S3.

### ΔΨ measurements

The ΔΨ of digitonin-permeabilized *T. brucei* cells was determined using the fluorescent dye Safranin O. 2x10^7^ cells were harvested by centrifugation (10 min, 1500 g) and washed in ANT buffer containing 8 mM KCl, 110 mM K-gluconate, 10 mM mannitol, 10 mM NaCl, 10 mM free acid HEPES, 10 mM K_2_HPO_4_, 0.015mM EGTA potassium salt, 0.5 mg/ml fatty acid-free bovine serum albumin, and 1.5 mM MgCl_2_ at pH 7.25. The cell pellet was resuspended in 2 ml of ANT buffer with addition of 5 µM Safranin O (Sigma, S2255) and 4 µM digitonin. Fluorescence was recorded in a Hitachi F-7100 spectrofluorometer (Hitachi High Technologies) at a 5-Hz acquisition rate, using 495 nm excitation and 585 nm emission wavelengths. ATP was added ad 1 mM to induce the membrane polarization. To assess the levels of ΔΨ generated, FCCP was titrated starting at 20 nM going up to 140 nM. Samples were measured at room temperature and stirred during the experiment.

### Alamar blue assays

Each cell line was inoculated in a transparent flat-bottomed 96-well plate at a density of 1×10^4^cells /ml in a total volume of 100µL of culture medium. Cells were grown at various concentrations of FCCP ranging from 0.025 µM to 166 µM and incubated at 37°C and 5% CO_2_ for 45 hours. Subsequently, 10 µl resazurin in PBS (0.5 mM) was added to each well and the cells were incubated for the next 3 hours. Fluorescence was then measured using the Tecan M1000 infinite pro plate reader at a wavelength of 536 nm for excitation and 588 nm for emission. The data was analyzed with GraphPad Prism 10.0 software using nonlinear regression and sigmoidal dose–response analysis.

### Correlation analysis: TbTim20-dependency versus N-terminal positive charges or pI

Protein sequences were retrieved from the TREU927 reference genome on TriTrypDB (65) and the pI and net charge used for Fig. S4 were calculated as follows. The isoelectric point (pI) of all quantified proteins predicted to have a mitochondrial presequence (n=94) (identified by TargetP) (44) was calculated in R (version 4.3.2) using the package Peptides (version 2.4.6) (function: pI, pKscale: EMBOSS) (66). The theoretical net charge of the first 20 aa of the same proteins was calculated scoring arginine and lysine residues as (+1) and aspartate and glutamate residues as (-1). (+1) was added to all final scores to take into account the protonation on the N-terminus. The final value corresponds to the theoretical net charge at approximately pH 7.4.

## Data availability

The mass spectrometry proteomcis data have been deposited to the ProteomeXchange Consortium (67) via the PRIDE (68) partner repository and are accessible using the dataset identifiers PXD061390 (TbTim20 CoIP data), PXD061391 (TbTim15 CoIP data), and PXD061395 (TbTim20 RNAi data).

## Acknowledgments

We thank Elke Horn and Thomas Morgenbrodt for excellent technical assistance, Pascal Mäser for help with the Alamar blue assays, Julian Bender for support in proteomics data analysis, and Christos Chinopoulos from Semmelweis University (Hungary) for helpful discussions. Research in the lab of BW was supported by the Deutsche Forschungsgemeinschaft (DFG, German Research Foundation) – SPP2453 Project ID 541758684. AZ was funded by OP JAK CZ.02.01.01/00/22_008/0004575 RNA for therapy, co-funded by the European Union and by the European Research Council project no. European Union and by the European Research Council. ASchneider was supported in part by NCCR RNA & Disease, a National Centre of Competence in Research (grant number 205601) and by project grant SNF 205200 both funded by the Swiss National Science Foundation. (69)

## Supplementary Figures

**Figure S1.**
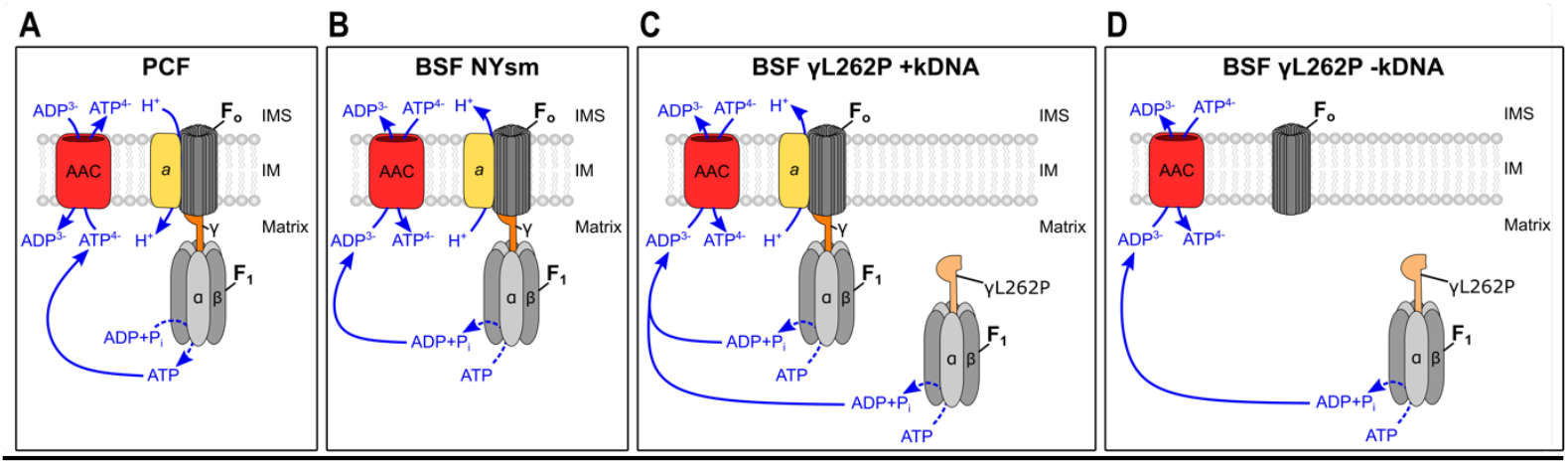
Mitochondrial ΔΨ generation in PCF and BSF trypanosomes. Suggested mode of ΔΨ generation in **(A)** PCF, **(B)** BSF NYsm, **(C)** BSF γL262P (+kDNA), and **(D)** BSF γL262P (+kDNA). ADP/ATP carrier (AAC, red), kDNA-encoded subunit *a* of the F_o_-ATPase (yellow), wild-type and mutant Ψ-subunits of the ATPase are shown in two shades of orange respectively. In (C) both wild-type and mutant version of the γ-subunit are present. It is unclear whether both versions can engage with both the coupled and uncoupled ATPase. It is not known in (C) and (D) whether the uncoupled F1-ATPase is completely detached from the IM. It is also unclear in (D) how much of the Fo-ATPase is present in the assembled state. IMS: intermembrane space, IM: inner mitochondrial membrane.

**Figure S2.**
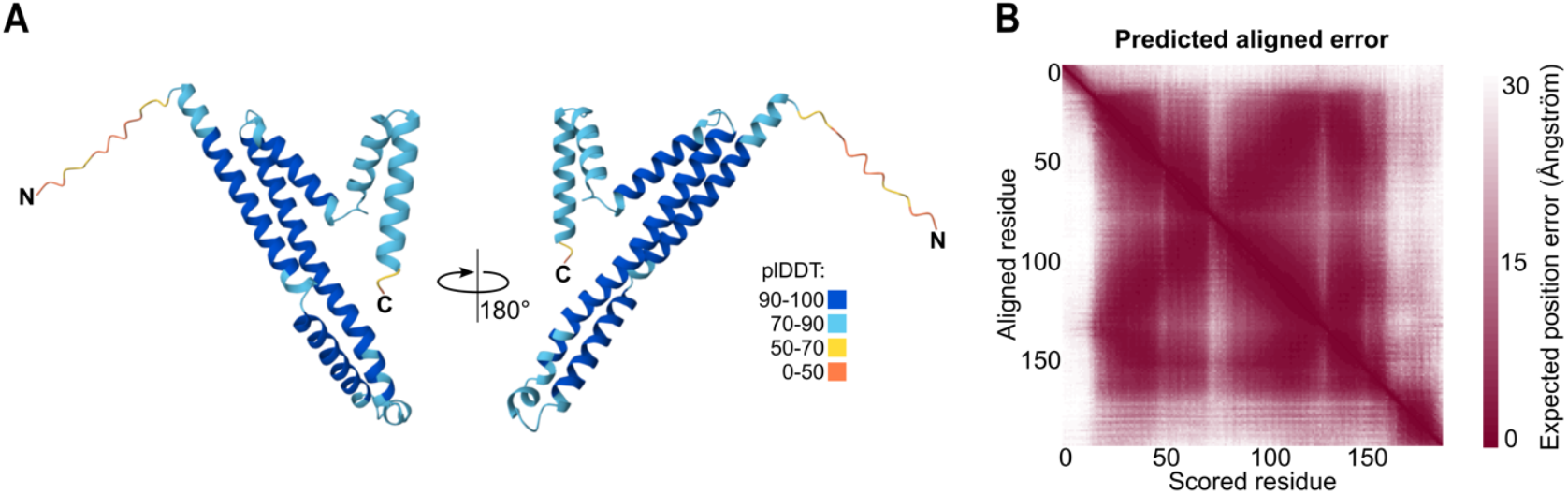
AlphaFold2 predicted structure of TbTim20. **(A)** Cartoon representation of the predicted monomer structure of TbTim20 from two sides. Colors represent the predicted local Distance Difference Test (plDDT). **(B)** Graphical visualization of the predicted alignment score in Angstrom of the scored residues versus the aligned residues. The model is available in ModelArchive (69) at https://www.modelarchive.org/doi/10.5452/ma-ow6rg.

**Figure S3.**
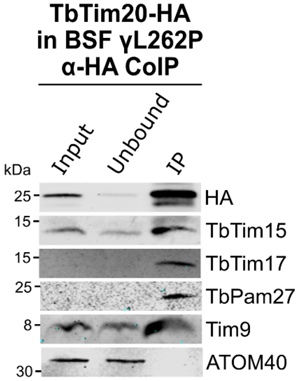
TbTim20 is a subunit of the TIM complex in γL262P cells. The γL262P(+kDNA) cell line expressing TbTim20-HA was subjected to a co-immunoprecipitation (CoIP) experiment. 5% of mitochondria-enriched fractions (Input), 5% of unbound proteins (Unbound) and 100% of the final eluates (IP) were analysed by immunoblotting. The immunoblot was probed with an anti-HA antibody, as well as with antisera recognizing the TIM/PAM subunits (TbTim15, TbTim17, TbPam27, Tim9) and ATOM40.

**Figure S4.**
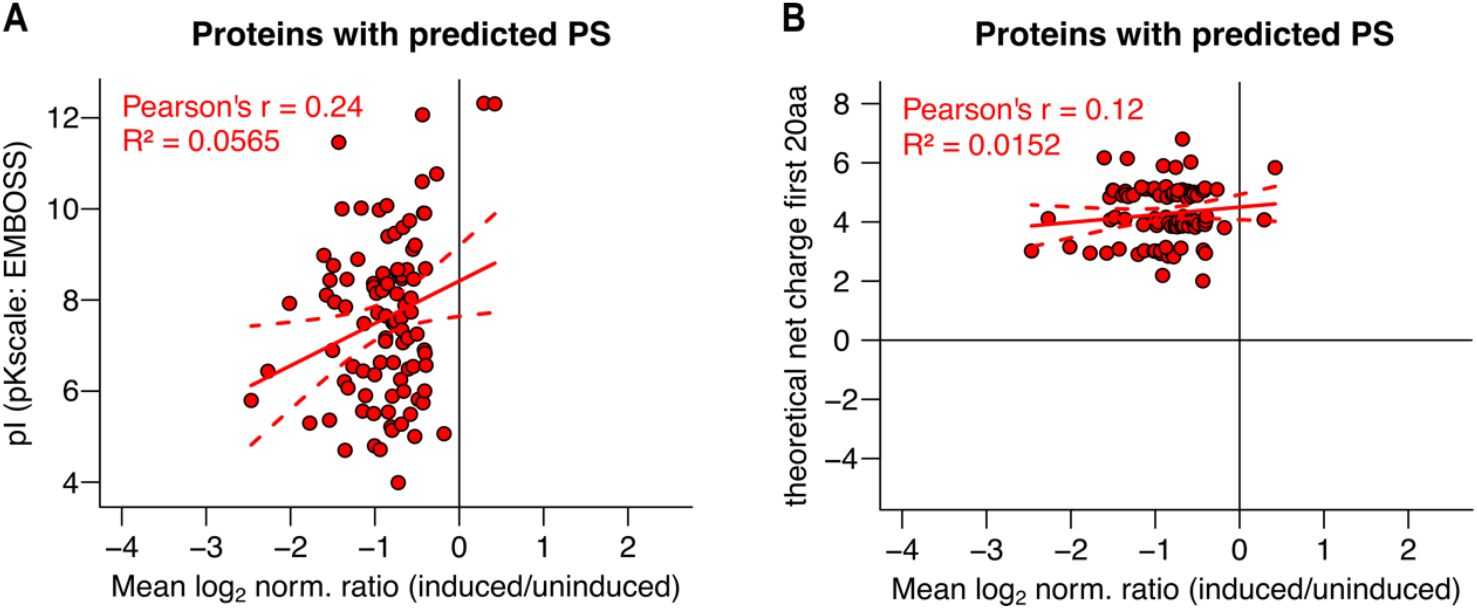
TbTim20-dependency not due to high N-terminal positive charges or pI. Graphs show the effect of the protein isoelectric point (pI) **(A)** or the theoretical net charge of the first 20 aa **(B)** on the mean log_2_ normalized ratio (induced/uninduced) of proteins with a predicted presequence (PS) (n=94). Pearson’s r: Pearson correlation coefficient; R^2^: Coefficient of determination for linear model. Solid lines show a linear regression line, dashed lines indicate the 90% confidence intervals of the linear regression. The values for the theoretical net charge of the first 20 aa in (B) were randomly scattered by maximal 0.4 units along the y-axis for better datapoint discrimination.

